# Predicting neoadjuvant treatment response in triple-negative breast cancer using machine learning

**DOI:** 10.1101/2023.04.17.536459

**Authors:** Shristi Bhattarai, Geetanjali Saini, Hongxiao Li, Hongyi Duanmu, Gaurav Seth, Timothy B. Fisher, Emiel A.M. Janssen, Umay Kiraz, Jun Kong, Ritu Aneja

## Abstract

**Background:** Neoadjuvant chemotherapy (NAC) is the standard treatment for early-stage triple negative breast cancer (TNBC). The primary endpoint of NAC is a pathological complete response (pCR). NAC results in pCR in only 30%–40% of TNBC patients. Tumor-infiltrating lymphocytes (TILs), Ki67 and phosphohistone H3 (pH3) are a few known biomarkers to predict NAC response. Currently, systematic evaluation of the combined value of these biomarkers in predicting NAC response is lacking. In this study, the predictive value of markers derived from H&E and IHC stained biopsy tissue was comprehensively evaluated using a supervised machine learning (ML)-based approach. Identifying predictive biomarkers could help guide therapeutic decisions by enabling precise stratification of TNBC patients into responders and partial or non-responders.

**Methods:** Serial sections from core needle biopsies (n=76) were stained with H&E, and immunohistochemically for the Ki67 and pH3 markers, followed by whole slide image (WSI) generation. The resulting WSI triplets were co-registered with H&E WSIs serving as the reference. Separate mask region-based CNN (MRCNN) models were trained with annotated H&E, Ki67 and pH3 images for detecting tumor cells, stromal and intratumoral TILs (sTILs and tTILs), Ki67^+^, and pH3^+^ cells. Top image patches with a high density of cells of interest were identified as hotspots. Best classifiers for NAC response prediction were identified by training multiple ML models, and evaluating their performance by accuracy, area under curve, and confusion matrix analyses.

**Results:** Highest prediction accuracy was achieved when hotspot regions were identified by tTIL counts and each hotspot was represented by measures of tTILs, sTILs, tumor cells, Ki67^+^, and pH3^+^ features. Regardless of the hotspot selection metric, a complementary use of multiple histological features (tTILs, sTILs) and molecular biomarkers (Ki67 and pH3) resulted in top ranked performance at the patient level.

**Conclusions:** Overall, our results emphasize that prediction models for NAC response should be based on biomarkers in combination rather than in isolation. Our study provides compelling evidence to support the use of ML-based models to predict NAC response in patients with TNBC.

## Introduction

Triple-negative breast cancer (TNBC) is an aggressive subtype of breast cancer (BC) and is often diagnosed at an advanced stage (1). TNBC is characterized by poor prognosis and high rates of recurrence and metastasis (2). No endocrine therapies are available for TNBC, and neoadjuvant chemotherapy (NAC) remains the mainstay of treatment for early-stage disease (3). A small subset of patients respond to newer therapies such as poly (ADP-ribose) polymerase inhibitors and immunotherapy (4). NAC helps to reduce tumor size before surgery (5), and its primary endpoint is a pathological complete response (pCR), defined as the absence of residual disease (RD). NAC results in pCR in only 30%–40% of TNBC patients, and the remaining patients either respond moderately or are refractory to NAC (i.e., show RD) (6). Conventional cytotoxic NAC for TNBC patients in the US consists of adriamycin, cyclophosphamide, and taxol (3). Although the molecular basis of chemoresistance in TNBC remains elusive, inter- and intra-tumoral heterogeneity may contribute to the significant variability in NAC response observed in patients with TNBC (7).

Existing biomarkers that predict NAC response in TNBC include Ki67 and phosphohistone H3 (pH3), which capture the proliferative potential and mitotic activity of tumor cells, respectively (8, 9). Although Ki67 scoring captures the proportion of cells that have entered the cell cycle (proliferative population or P), it may not accurately represent proliferation because a Ki67-positive cell (Ki67^+^) may not divide for long periods of time. Histone H3 is heavily phosphorylated (pH3-positive) during mitosis (10); therefore, pH3 staining can capture true proliferation based on the identification of actively dividing cells (pH3^+^/mitotic population/M). Cells stained positive for pH3 are considered to be Ki67-positive (Ki67^+^/pH3^+^), whereas tumor cells positive for Ki67 may (Ki67^+^/pH3^+^) or may not (Ki67^+^/pH3^−^) be pH3-positive, depending on their mitotic status. Tumor assessment using hematoxylin and eosin (H&E)-stained slides of surgical resections or biopsies provides additional histological information that can predict pCR, such as the number of tumor-infiltrating lymphocytes (TILs) (11-14). However, systematic evaluation of the ability of these biomarkers to predict NAC response has not yet been conducted.

TNBCs are complex evolving systems characterized by profound spatial and temporal heterogeneity in their biological nature and response to treatment. Individual biomarkers that depict only a single aspect of tumor physiology or biophysics are limited, and their predictive performance may vary among tumors. A multiparametric approach that combines information from functional imaging technologies with complementary sensitivity is required to predict NAC outcomes in patients with TNBC. In this study, we developed a machine learning (ML) approach that integrates Ki67, pH3 and TILs [both intratumoral (tTILs) and stromal (sTILs)] in co-registered serial whole slide images (WSIs) to predict NAC response in patients with TNBC.

## Materials and methods

### Study cohort

A total of 76 formalin-fixed paraffin-embedded (FFPE) TNBC biopsy samples were retrieved from Emory Decatur Hospital. Biopsy samples were collected before systemic treatment. Of these patients, 44 showed pCR and the remaining 32 showed RD after NAC treatment. All study aspects, including study protocols, sample procurement, and study design, were approved by the Institutional Review Board (IRB). Clinicopathological data, patient survival information, and NAC response data were available. For each patient, three tissue slides containing serial sections (5 μm) were stained with H&E, Ki67, and pH3.

### Serial image co-registration

High-resolution WSIs of three serial tissue slides were produced for each patient; one was stained with H&E, and two were immunohistochemically stained for Ki67 and pH3. WSI triplets were co-registered at the highest image resolution using our previously developed dynamic co-registration method (15). For each image triplet, the H&E WSI served as the reference image and the other two IHC images were mapped to the reference image. After image registration, our dataset consisted of 1,044 WSI image regions of 8,000×8,000 pixels from 76 TNBC patients. As the registered images were large in size and Graphical Processing Units (GPUs) for model training and testing have limited memory, each image region was further partitioned into non-overlapping image patches of 1,000×1,000 pixels by size. The resulting image patches were appropriate for the ML analyses.

### H&E staining

FFPE samples were subjected to a series of xylene washes followed by alcohol washes. The tissues were then thoroughly rinsed with water and stained with Harris hematoxylin. Subsequently, tissue sections were stained with eosin, which stains nonnuclear elements in different shades of pink. After rinsing in a series of alcohol solutions and xylene, a thin layer of polystyrene mountant was applied, followed by tissue mounting on a glass coverslip.

### Immunohistochemistry

Immunohistochemistry (IHC) was performed as previously described (15). FFPE tissue sections were deparaffinized by a 20-minute incubation in an oven, followed by a series of xylene washes. The tissues were rehydrated in a series of ethanol solutions (100%, 90%, 75%, and 50%). Antigen retrieval was performed by heating tissues in citrate buffer (pH 6.0) using a pressure cooker (15 psi) for 30 min. The tissues were cooled to room temperature and then incubated in hydrogen peroxide for 10 min, followed by blocking in UltraVision Protein Block (Life Sciences Inc., FL, USA) for 10 min. Tissue samples were incubated for 60 min at room temperature with primary antibodies (Monoclonal Mouse Anti-Human Ki67, clone MIB-1, Dako North America Inc., FL, USA at 1:100 dilution; Phosphohistone H3 (pH3), Biocare, 1:500 dilution). After a series of washes, tissues were incubated with a MACH2 HRP-conjugated secondary antibody (Biocare Medical, CA, USA). Enzymatic antibody detection was performed using Betazoid DAB Chromogen Kit (Biocare Medical, CA, USA). Finally, the tissue sections were counterstained with Mayer’s hematoxylin, dehydrated in a series of ethanol concentrations, and mounted with mounting media.

### Imaging and tumor slide annotation

Slides were scanned using a slide scanner (Hamamatsu NanoZoomer 2.0-HT C9600-13) at 40x magnification (0.23 μm/pixel). Using Aperio ImageScope 12.4.3, a board-certified pathologist reviewed the images for quality, overlapping tissue areas, out-of-focus areas, and staining artifacts. Tumor cells, sTILs and tTILs were manually annotated in H&E WSIs. According to international guidelines (16), tTILs were defined as lymphocytes in a tumor cell nest in direct contact with adjacent tumor cells, and sTILs were defined as lymphocytes in the tumor stroma and not in direct contact with tumor cells. Artifacts and necrotic areas were manually excluded from evaluation.

### ML pipeline

To emulate a pathologist’s review process, we developed a traditional ML prediction pipeline (**Figure 1**) consisting of multiple processing steps, including serial histopathological image generation (**Suppl Figure 1**), registration, cell detection, biomarker identification, hotspot region detection, hotspot feature extraction, classifier optimization, and NAC response prediction.

**Figure 1.**
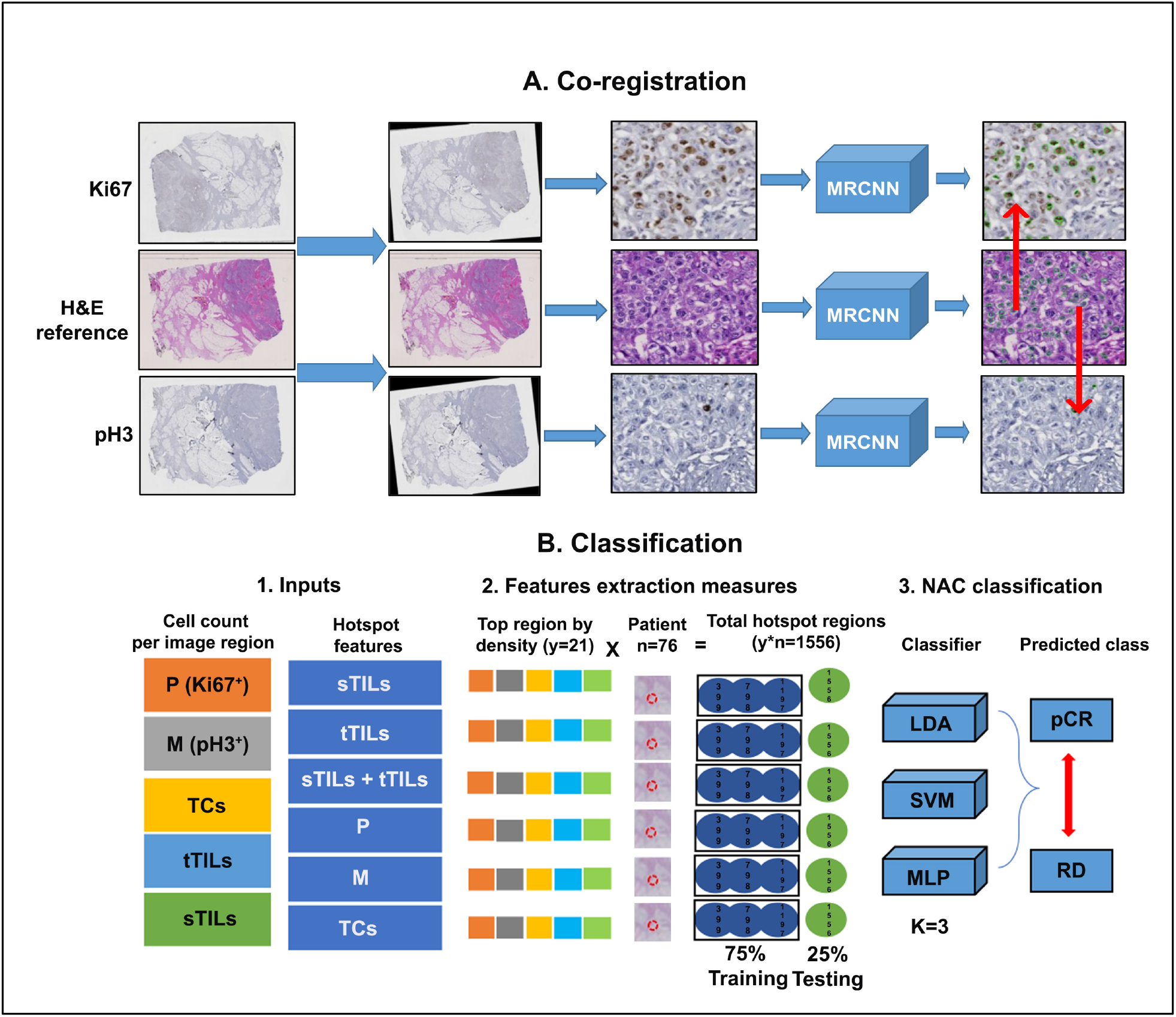
Schema of our ML pipeline. A) Co-registration-Three WSIs (H&E, IHC-Ki67, and IHC-pH3) for each patient were acquired and spatially registered at the highest image resolution before being partitioned into image patches of 1,000 × 1,000 pixels. Multiple MRCNN models were trained and applied to image patches to detect TCs (tumor cells), TILs (tumor infiltrating lymphocytes) in H&E slides, P (tumor cells stained positive for Ki67) and M (tumor cells stained positive for pH3) in IHC images. B) Classification-This part of the pipeline includes inputs of image patches and cell counts. The image patches with the highest spatial density of cells of interest were identified as diagnostic hotspots. The number of TCs, P, M, sTILs, and tTILs from the hotspots were used as hotspot features to train (75%) and test (25%) the classification models for NAC response prediction.

#### Cell detection and phenotyping

Tumor cells, TILs, and cells stained with IHC biomarkers in spatially aligned WSIs were detected using Mask R-CNN (MRCNN) (17) trained with H&E, Ki67, and pH3 image patches. The MRCNN model for tumor cell detection was trained using 797 H&E images of 1,000 × 1,000 pixels with all tumor cells annotated by multiple expert human reviewers. The MRCNN model for TIL detection was trained using 500 H&E images of 1,000 × 1,000 pixels with TIL annotations by pathologists. In contrast, the MRCNN models for Ki67^+^ and pH3^+^ tumor cell detection were trained using 30 and 20 well-annotated IHC images (1,000 × 1,000 pixels), respectively. Cells of interest for NAC response prediction included tumor cells, sTILs, tTILs, Ki67^+^, and pH3^+^. sTILs and tTILs were marked by pathologists, whereas Ki67^+^ and pH3^+^ were identified by mapping serial IHC images to the reference H&E image (Figure. 1A). Tumor cells in H&E images were identified as Ki67^+^ or pH3^+^ if Ki67 or pH3 staining was detected within a radius of 40 pixels. This radius is empirically determined based on the average tumor cell size.

#### Identification of hotspot regions

We identified hotspot regions enriched with cells of interest in WSIs. Cell types of interest for hotspot recognition included sTILs, tTILs, sTILs + tTILs, Ki67^+^ (henceforth referred to as P), pH3^+^ (henceforth referred to as M), and tumor cells (TCs). For each selection metric, all image patches from each patient were sorted in descending order, and the top image patches with a high cell density were selected as hotspots. A typical hotspot selected by pathologists includes 500–2000 tumor cells (18, 19). Given that each image patch in our dataset included an average of 100 tumor cells, we included a sufficiently large number of tumor cells by considering the top 21 image patches as hotspots. The odd hotspot number made it easier to support majority voting while deriving the predicted patient class label from the hotspot labels.

#### Classifier optimization and NAC response prediction

The NAC response predictor was trained using cell profile features extracted from the hotspot regions. Features derived from each hotspot included counts of sTILs, tTILs, M, P, and TCs. All hotspots from the same patient shared the same patient-class label (i.e., either pCR or RD). To identify the best classifiers for NAC response prediction, we used multiple ML methods, including linear discriminant analysis (LDA), support vector machine (SVM), and multilayer perceptron (MLP). We optimized the hyperparameters of these ML algorithms using the auto-ML tool Auto-Sklearn (20). The resulting hotspot dataset included 1,596 image patches from 76 patients (44 with pCR, and 32 with RD) and was randomly split into 75% for training (57 patients; 33 with pCR, and 24 with RD) and 25% for testing (19 patients; 11 with pCR, and 8 with RD). Prediction performance was evaluated using a three-fold cross-validation method. The trained model with the highest average validation accuracy was retained and applied to the testing data. For analysis pipeline implementation, pCR was defined as the positive class and RD as the negative class.

Prediction accuracy was defined as:

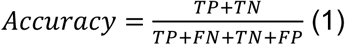

where TP, FP, TN, and FN represent true positive, false positive, true negative, and false negative, respectively.

The true positive rate (TPR) and the false positive rate (FPR) were defined as:

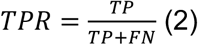

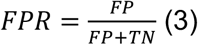

Additionally, prediction accuracy was evaluated by the area under the receiver operating characteristic curve (ROC AUC). ROC analysis was performed to assess the diagnostic ability of the classifiers, by plotting the true positive rate against the false positive rate. The ROC AUC reflects the ability of the classifier to distinguish between positive and negative cases (21).

## Results

### Prediction accuracy of ML classifiers for different hotspots

The prediction accuracies of the optimal ML classifiers for hotspots by different selection creiteria (i.e., tTILs, sTILs, sTILs + tTILs, M, P, TCs) are presented in **Figure 2** where each hotspot is represented by tTILs, sTILs, P, M, and TCs. At the patient level, the highest prediction accuracy was achieved with the tTILs classifiers. In contrast, patient-level prediction accuracy for sTILs hotspots was relatively low. When sTILs and tTILs were jointly considered (sTILs + tTILs) as hotspots, the resulting prediction performance lay between the prediction accuracies of the individual hotspots. In general, the prediction accuracy at the image patch level was lower than that at the patient level. Because of the prediction robustness introduced by the maximum voting mechanism, the patient-level prediction performance was not significantly affected by the misclassification rates at the image patch level. The patient-level prediction accuracy for M or P hotspots was lower than that for the TILs hotspots. Although the patient-level prediction accuracy was higher for P hotspots than for M hotspots, the opposite trend was observed for the image patch-level prediction accuracy (**Figure 2**).

**Figure 2.**
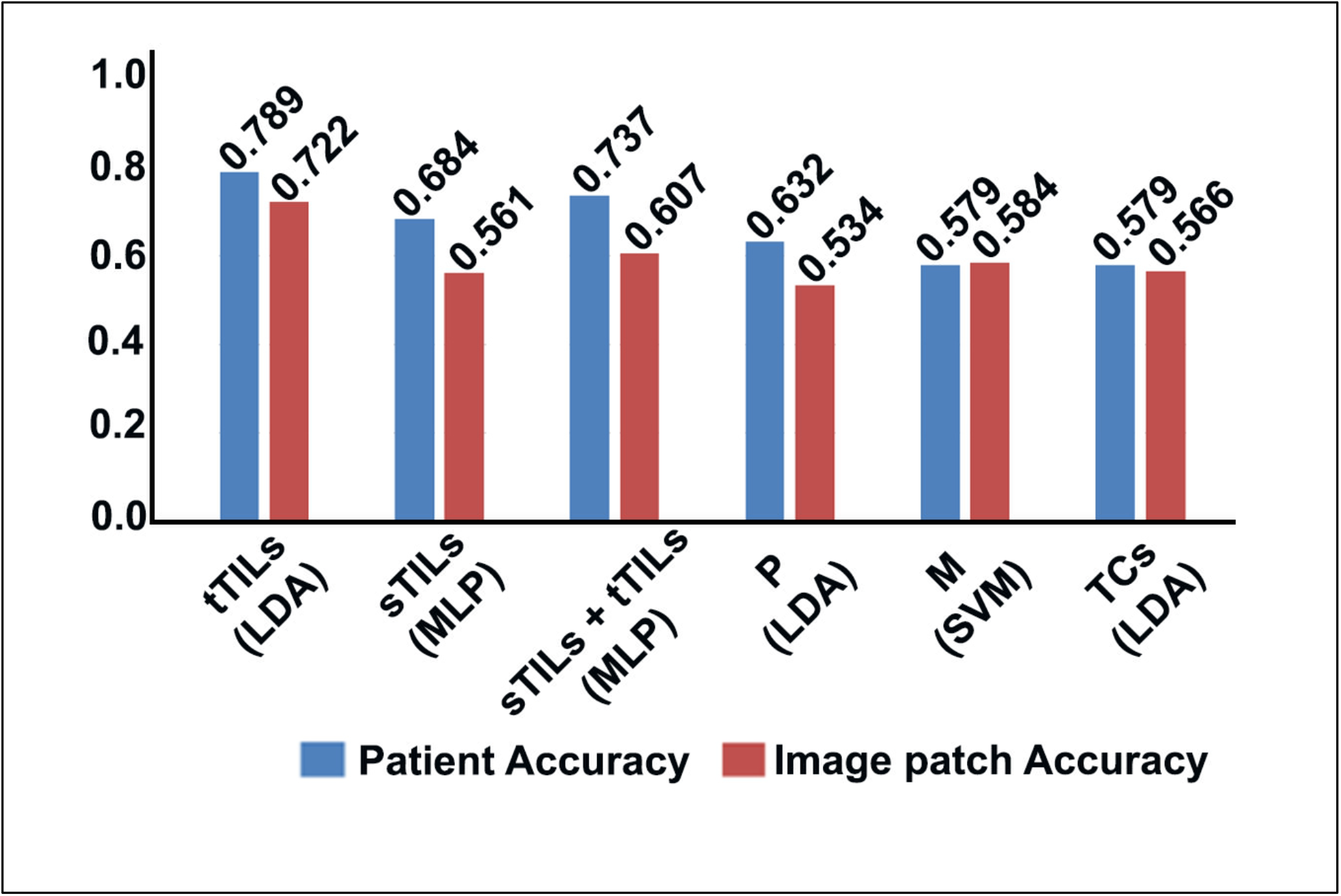
Prediction accuracies of the optimal ML classifiers using different hotspot selection standards with hotspot feature representation by tTILs, sTILs, P, M, and TCs. In contrast, MLP showed the best performance when the count of sTILs or sTILs + tTILs was used as hotspot selection standard. SVM performed best when M was used as the hotspot selection standard. LDA = linear discriminant analysis; SVM = support vector machine; MLP = multilayer perceptron

Receiver operating characteristic (ROC) curve analysis was conducted to calculate the area under the curve (AUC) values for the different prediction models (**Figure 3**). Prediction performance for tTILs hotspots was the highest among all hotspots, demonstrating the considerable impact of hotspot selection standard on the prediction results. When tTILs and sTILs were jointly considered for hotspot selection, the AUC value was higher than the AUC value obtained with the model for sTILs hotspots prediction (**Figure 3**). Moreover, the AUC value for M hotspots was higher than that for P hotspots, but lower than the TCs hotspots.

**Figure 3.**
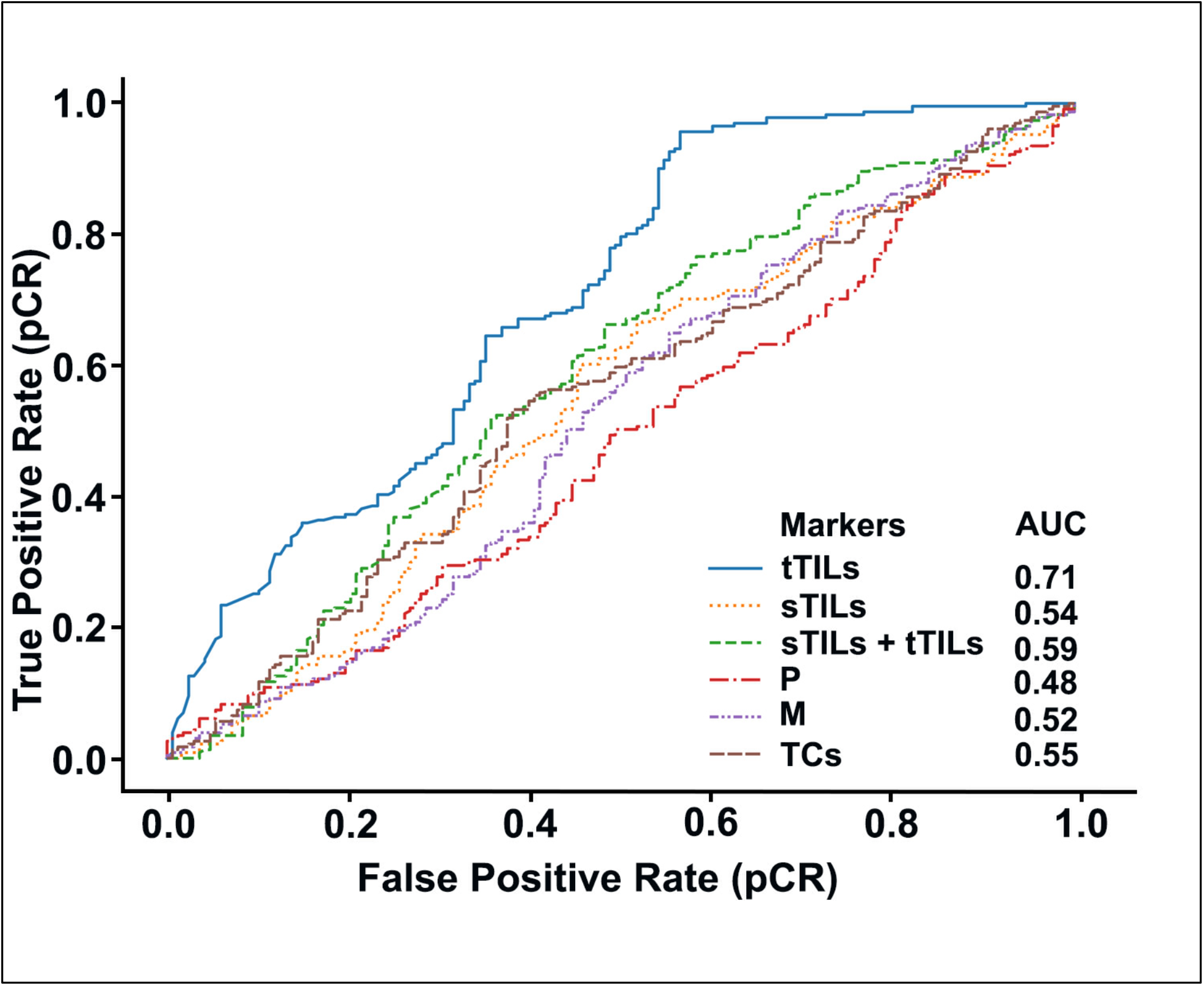
ROC curves of the best classifiers according to different hotspot selection standards. Best classifier performance was observed in tTILs (AUC = 0.71), followed by sTILs + tTILs (AUC = 0.59).

### Biomarker hotspots accurately predict pCR in patients with TNBC

The patient- and image patch-level testing performances of the optimal prediction models for different hotspots were compared using confusion matrix analysis (**Figures 4 and 5**). Regardless of the hotspot standard, all prediction models correctly identified all patients with pCR (except the sTILs hotspots model that misclassified one case) with no false-negative results at the patient level. However, different hotspot detection models showed variable RD prediction performance. The misclassification rate for RD varied from 50% with tTILs hotspots to 100% with M or TC hotspots. Similar results were obtained at the image patch level (**Figure 5**). Importantly, the image patch-level prediction accuracy for pCR reached 92% with the tTILs hotspots.

**Figure 4.**
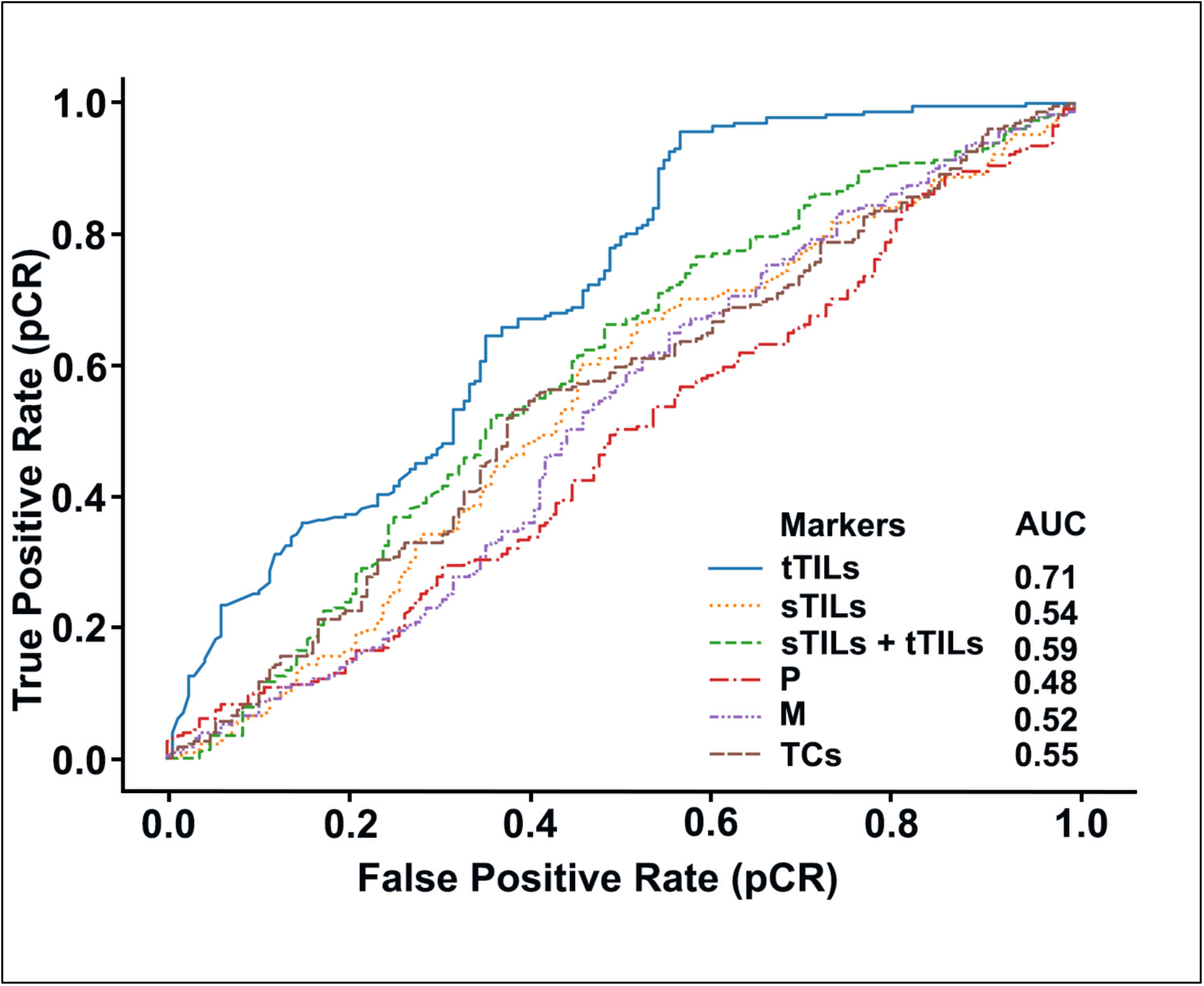
Confusion matrices showing patient-level testing performance. When hotspots were identified by the count of tTILs, sTILs, sTILs + tTILs, P, M, and TCs, the associated best classifiers showed different performances; pCR prediction performance was accurate, with variable RD prediction performance. The patient-level prediction label was determined using patch-level labels by maximum voting. TP (true positive); TN (true negative); FP (false positive); FN (false negative).

**Figure 5.**
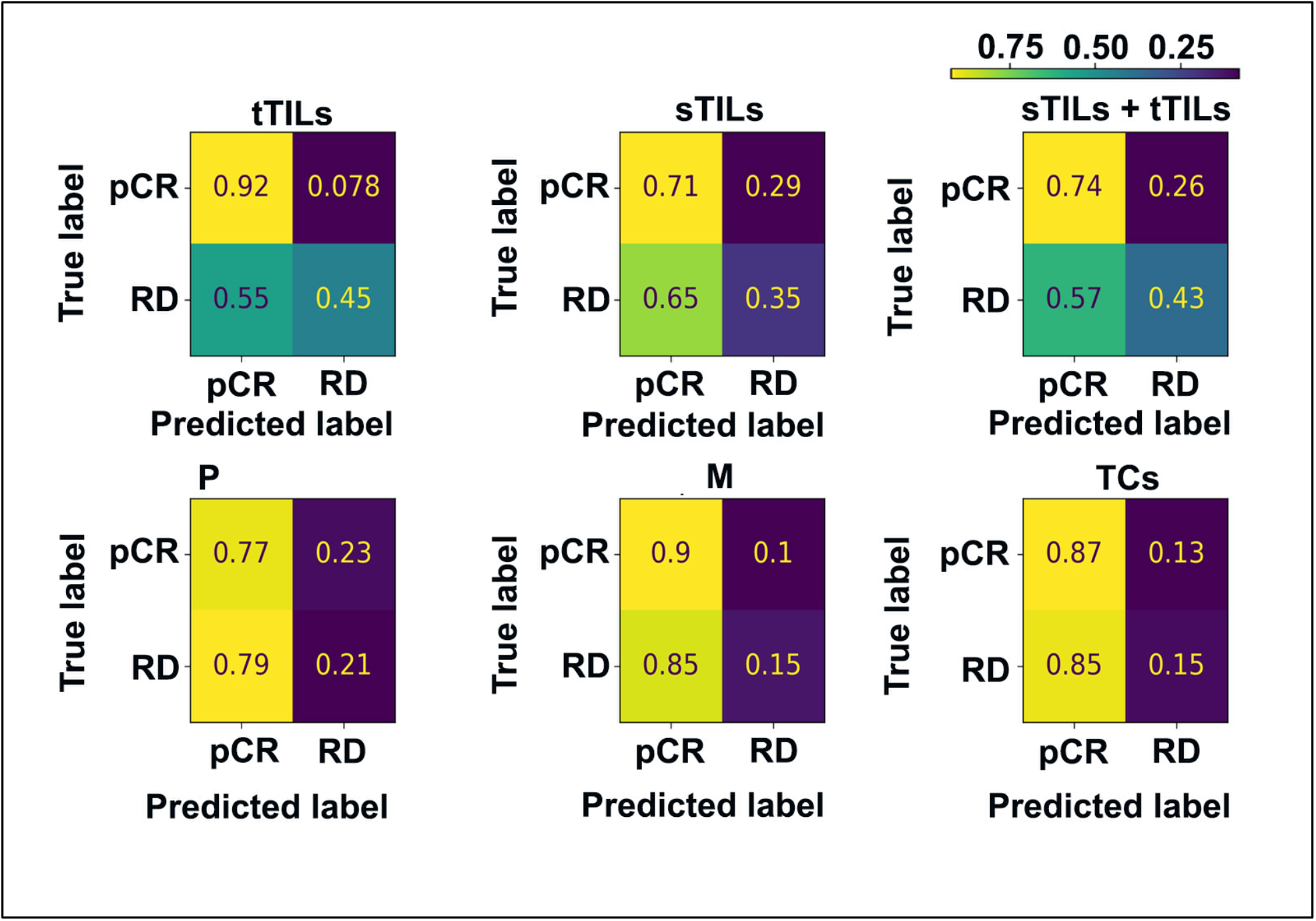
Confusion matrices showing image patch-level testing performance. When the hotspots were identified by the count of tTILs, sTILs, sTILs + tTILs, P, M, and TCs, the associated best classifiers showed different performances as suggested by the patch-level confusion matrices with values normalized by rows. While the pCR prediction accuracy was high (92% with tTILs hotspots), the RD prediction accuracy was variable.

### Prediction accuracy associated with different hotspot features

We investigated the best prediction accuracies for hotspot features at the image patch and patient levels using different hotspot selection metrics (**Table 1**). Overall, hotspots identified by tTIL counts (i.e., tTILs hotspots) showed the best prediction accuracy at the patient level. The combination of the tTIL count and additional cell counts (sTILs, P, and M) enhanced the prediction accuracy at both the image patch and patient level. Similarly, for the other hotspot selection metrics, the use of multiple cellular features (i.e., counts of tTILs, sTILs, and TCs) and molecular biomarkers (i.e., P and M) resulted in improved prediction performance, suggesting the presence of complementary prediction values from these classes of hotspot features (H&E and IHC derived data).

**Table 1.** The Accuracies of hotspots with various feature combinations Footnote: TCs = Tumor Cells; tTILs = intratumoral TILs; sTILs = stromal TILs; P = Ki67^+^; M = pH3^+^; LDA = linear discriminant analysis; SVM = support vector machine; MLP = multilayer perceptron; ML = machine learning

| Hotspot identification metric | Features considered in hotspots | Image patch accuracy | Patient accuracy | ML algorithm |
| --- | --- | --- | --- | --- |
| **TCs** | TCs | 0.579 | 0.579 | LDA |
|  | tTILs; sTILs; P; M; TCs | 0.566 | 0.579 | LDA |
|  | tTILs; sTILs; P; M | 0.576 | 0.579 | SVM |
| **tTILs** | tTILs | 0.702 | 0.684 | LDA |
|  | tTILs; sTILs; P; M ; TCs | 0.722 | 0.789 | LDA |
|  | tTILs; sTILs; P; M | 0.739 | 0.789 | LDA |
| **sTILs** | sTILs | 0.634 | 0.632 | LDA |
|  | tTILs; sTILs; P; M; TCs | 0.561 | 0.684 | MLP |
|  | tTILs; sTILs; P; M | 0.632 | 0.632 | LDA |
| **sTILs + tTILs** | tTILs; sTILs | 0.657 | 0.632 | LDA |
|  | tTILs; sTILs; P; M; TCs | 0.607 | 0.737 | MLP |
|  | tTILs; sTILs; P; M | 0.632 | 0.632 | LDA |
| **P** | P | 0.534 | 0.526 | MLP |
|  | tTILs; sTILs; P; M; TCs | 0.534 | 0.632 | LDA |
|  | tTILs; sTILs; P; M | 0.574 | 0.526 | LDA |
| **M** | M | 0.499 | 0.421 | LDA |
|  | tTILs; sTILs; P; M; TCs | 0.584 | 0.579 | SVM |
|  | tTILs; sTILs; P; M | 0.599 | 0.579 | LDA |

## Discussion

pCR is an important predictor of improved disease-free survival (DFS) and overall survival (OS) in patients with TNBC (22). pCR has also been associated with favorable prognosis in neoadjuvant trials in TNBC and has become a surrogate marker of survival (22). Existing markers of NAC response, i.e., predictive of pCR, include Ki67, pH3, and TILs. However, there are no established biomarkers of NAC response using diagnostic TNBC biopsies that can be readily adopted in clinical practice. Additionally, there is a lack of comprehensive assessment of the predictive value of potential biomarkers for NAC response in patients with TNBC. This lack of readily quantifiable predictive biomarkers that can accurately risk-stratify patients with TNBC is a barrier to optimal therapeutic decision-making. Identifying predictive biomarkers could help guide therapeutic decisions by enabling precise stratification of TNBC patients into responders and partial or non-responders, who may be spared the unnecessary adverse effects of NAC, and directly receive surgery. Predictive biomarkers could also help clinicians identify immunomodulatory strategies for partial responders, which, in combination with chemotherapy, may enhance the response to NAC and improve treatment outcomes. In this study, we have comprehensively evaluated the predictive value of markers derived from H&E and IHC stained biopsy tissue, using a supervised ML-based approach. We analyzed Ki67, pH3, and TILs levels in serial WSIs. We also spatially integrated biomarkers and histological features to predict NAC response. The spatial relationship across serial WSIs was established using image co-registration (15). ML algorithms were applied to these spatially aligned images to detect Ki67, pH3, and TIL hotspots. ML methods can outperform human reviewers in terms of accuracy, speed, and reproducibility (23). Additionally, they can reveal predictive morphological features and spatial patterns beyond human perceptual abilities (24, 25).

Our findings suggest that the hotspot selection metric significantly affects the prediction performance of the model. Notably, TILs (derived from H&E-stained tissue slides) exhibited strong predictive accuracy, outperforming P (Ki67^+^, derived from IHC stained tissue sides), which is the current clinical standard. Our data suggest that the tTILs count is the optimal standard for hotspot selection. H&E staining is the gold standard for histopathological assessment of the tumor microenvironment (TME) (26). H&E-stained WSIs provide vital information about tissue architecture, including the type of cells (e.g., epithelial, stromal, and TILs) and their spatial arrangement in the dynamic TME. Several studies have shown that histopathological TME components, particularly TILs, can be used to predict pCR because increased counts of sTILs and tTILs have been correlated with pCR in TNBC patients (27, 28). TILs mirror the local immune response, and high TIL counts are associated with favorable OS in patients with TNBC (29-31). A better understanding of the immune landscape can uncover the connections between immune components and therapeutic response, yielding novel predictive markers for NAC response in TNBC.

After hotspot identification, we used different combinations of cellular features (counts of tTILs, sTILs, and TCs) and molecular biomarkers (Ki67 and pH3) to predict NAC response. The counts of sTILs, tTILs, P, and M, considered in tTIL hotspots, exhibited the highest predictive value. The combination of histological features from H&E images and molecular biomarkers from IHC images performed better than individual features, supporting the complementary predictive ability of these two classes of features. Overall, our results emphasize that prediction models for NAC response should be based on biomarkers in combination rather than in isolation.

Tumors with a higher level of cell proliferation respond better to chemotherapy than tumors with a lower level of proliferation (32, 33). Ki67 index is a routinely used clinical marker of cancer cell proliferation and is strongly correlated with recurrence and metastasis (34-36). Nuclear Ki67 is expressed in all active phases of the cell cycle (G1, S, and G2) (37, 38). Patients with high Ki67 levels respond well to NAC (39), and thus, Ki67 levels can independently predict pCR. However, the cutoff value for classifying Ki67 expression as high or low is not standardized and ranges from 12% to 25% (40). Ki67 may fail to accurately represent actively dividing cells. Although Ki67 scoring captures the proportion of cells that have entered the cell cycle, it may misrepresent proliferation in the truest sense, as Ki67^+^ cells (proliferating or P population) may not divide for long periods of time, and cells in the G1 phase have uncertain fates (41-44). Current clinical Ki67 evaluation is highly subjective, causing differing opinions among pathologists while selecting fields for assessment of heterogeneous tumors, such as TNBC. Chemotherapy targets actively dividing cells during mitosis; hence, pH3, a marker of mitotic activity, has emerged as a predictor of NAC response. Histone H3 is a core histone protein, and its phosphorylation occurs exclusively during mitosis (S-phase) (10, 45); therefore, pH3 staining can capture true proliferation based on the identification of mitotic cells (M population). Although pH3 staining has not yet been clinically adopted, its ability to predict pCR has been demonstrated in several clinical studies and is superior to Ki67 staining in terms of reproducibility and ability to represent proliferation (10). In current practice, P (Ki67^+^) and M (pH3^+^) population of tumor cells are evaluated independently from a limited number of high-power tumor fields (46) and the predictive power of either variable alone may result in inaccurate patient risk stratification or clinical decision-making. Therefore, Ki67 should be considered in conjunction with pH3, in the same microscopic field or region of the tumor, to obtain a true picture of tumor cell proliferation. Traditional scoring methods for Ki67 and pH3 are time-consuming and prone to inter- and intra-observer variability, limiting the clinical value of these biomarkers. By contrast, ML methods can process entire slides for multiple markers concurrently, yielding enriched information in a fraction of the time. ML is less susceptible to inter- and intra-observer subjectivity than manual scoring and can identify novel predictive features and spatial patterns. Thus, ML-based prediction models present immense clinical potential and are poised to become mainstream tools for breast cancer diagnosis. By leveraging co-registered H&E and IHC WSIs of serial tissue sections, our ML-based pipeline integrates complementary information from histological components (TCs, tTILs, and sTILs in H&E slides) and biomarkers (Ki67 and pH3 in serial IHC slides) to predict NAC response. All components in this computational pipeline are fully automated, making it more efficient than traditional manual reviewing. Our findings suggest that the prediction accuracy from the feature P alone, considered within the P hotspot identification metric, is better than that from the feature M alone within the M hotspot identification metric, both at the image patch and patient level. Furthermore, a joint consideration of the features – sTILs, tTILs, M, P, and TCs-resulted in improved prediction accuracy at the image patch and patient levels, for the M and P hotspot identification metrics. Thus, considering both set of populations, actively dividing and the subset that is not, may yield improved predictive value.

This study has some limitations. In our pipeline, we considered select TME components (tumor cells, tTILs, sTILs,) and biomarkers (Ki67 and pH3). However, the pipeline can be extended to include a larger number of predictive biomarkers and clinicopathological data (e.g., age, tumor grade, stage, and lymph node status) to enhance the predictive ability of the model. Additionally, hotspots can be detected using different metrics and fixed cutoff values rather than using dynamic cutoffs determined by the data. These modifications may better mirror current clinical practice protocols and enhance the interpretability of NAC response prediction. Furthermore, the image patch-level results are aggregated with the patient-level results using the maximum voting rule. By doing so, we implicitly assume the equal impact of all image patches on the patient-level prediction results. We plan to optimize this aggregation rule at a better resolution in future research. Due to limitations of immunohistochemistry, a subset of M population may not be positively stained for Ki67 (appearing erroneously as pH3^+^/Ki67^−^), and similarly, a subset of P population that are in the mitotic cycle (should be Ki67^+^/pH3^+^) may not be positively stained for pH3 (appearing as Ki67^+^/pH3^−^). Imprecise co-registration can also result in a subpopulation of M cells lacking a Ki67signal (pH3^+^/Ki67^−^).

## Conclusion

Our study provides compelling evidence to support the use of ML-based models to predict NAC response in patients with TNBC. With its effectiveness and interpretability, our prediction model shows a promising potential for clinical adoption in future.

## List of abbrevations

## Declarations

### Ethics approval and consent to participate

All aspects of this study were approved by the Institutional Review Boards of the institutions involved. Patient consent was not required because all samples were archival.

### Consent for publication

NA

### Availability of data and materials

The data underlying this article will be shared on request to the corresponding author.

### Competing interest

The authors declare no conflicts of interest or financial interest.

### Funding

This study was supported by a National Cancer Institutes grant (R01CA239120) to RA and a National Institutes of Health grant (1U01CA242936) to JK .

## Authors Contribution

SB, GS and RA: Conceptualization

SB, GS: Data procurement, experimentation JK, HL, HD, TF: Data analysis

SB, GS, JK: Manuscript writing and editing

SB, GS, JK,EJ, UK and RA: Discussion, editing and proof-reading the manuscript

## Acknowledgments

Editorial support for this manuscript was provided by Christos Evangelou, Ph.D. We would like to thank Mahak Bhargava for managing the WSIs and all the institutions and hospitals involved in this study for providing us with tissue samples.

## Supplementary Files

**Figure S1.**
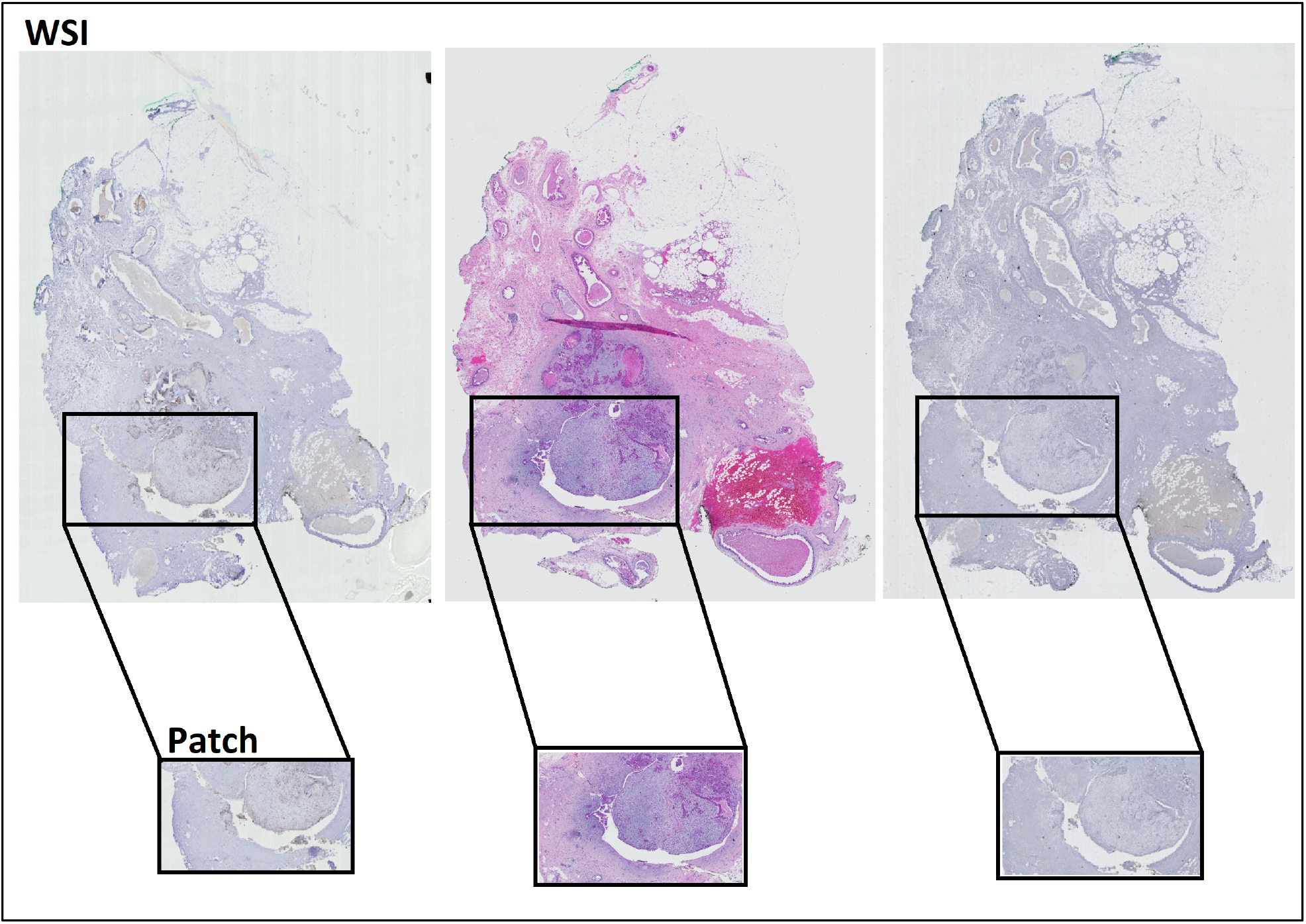
Typical example WSI triplets of a patient from our dataset. The WSI is the gigapixel image the represents the entire patient tissue biopsy. WSI triplets are registered by 8,000 × 8,000 regions that are further partitioned to 1,000 × 1,000 non-overlapping patches to facilitate the computation.

## Notes

### Competing Interest Statement

The authors have declared no competing interest.

